# HPV16 recruitment of SMARCAL1 to viral and host replication forks is required for the viral life cycle

**DOI:** 10.1101/2025.07.22.666081

**Authors:** Claire D James, Aya H Youssef, Jenny D Roe, Floriana Cappiello, Francesca Antonella Aiello, Benedetta Perdichizzi, Rachel L Lewis, Austin Witt, Apurva T. Prabhakar, Xu Wang, Molly L Bristol, Pietro Pichierri, Iain M Morgan

## Abstract

High-risk human papillomaviruses (HR HPVs) are responsible for around 5% of the world’s cancer burden. Activation and interaction with the host DNA damage response (DDR) promotes the HPV16 life cycle. This study demonstrates a crucial interaction between HPV16 and SMARCAL1, a protein involved in the stabilization of stalled DNA replication forks. SMARCAL1 can complex with E2, is recruited to E1-E2 replicating DNA, and SMARCAL1 knockdown reduces the fidelity of E1-E2 mediated DNA replication in C33a cells but does not alter replication levels. SMARCAL1 is recruited to the HPV16 genome in HPV16-immortalized foreskin keratinocytes (HFK+HPV16), and *in situ* protein interaction with nascent DNA replication forks (SIRF) assays demonstrated that SMARCAL1 is hyper-recruited to host replication forks in HFK+HPV16 cells. Using COMET, DNA fiber, and cell growth assays it was determined that knockdown of SMARCAL1 increased DNA damage and impaired replication fork progression in HFK+HPV16 cells, ultimately resulting in growth arrest. The viral genome integrates following SMARCAL1 knockdown in HFK+HPV16 cells. Therefore, SMARCAL1 facilitates host and viral DNA replication in HFK+HPV16 cells. Overall, the results demonstrate that HPV16 promotes SMARCAL1 recruitment to viral and host replication forks and is an essential factor for the HPV16 life cycle. The results expand our understanding of DDR proteins that regulate the HPV16 life cycle, and suggest that inhibition of SMARCAL1 function represents a novel anti-viral strategy for the treatment and prevention of HPV infections.

**Importance:** HPV16 is responsible for the majority of HPV+ cancers, contributing to 54% of cervical cancers and ∼90% of HPV+HNSCC. Integration of viral genomes into host DNA can promote cervical cancer progression and correlates with poor prognosis in HPV-associated HNSCC, where around 70% of HPV+ cancers contain episomal viral genomes. Developing effective antiviral therapies requires a deeper understanding of the interplay between viral replication and host DNA damage response (DDR) pathways. This report demonstrates that SMARCAL1 is essential for HPV16 replication and keratinocyte proliferation and that its depletion leads to replication stress, DNA damage, and viral genome integration. This work underscores the delicate balance between viral exploitation of the host DDR and the risk of genome instability. These insights contribute to the broader understanding of HPV pathogenesis and may inform the development of therapeutic strategies targeting viral replication to prevent disease progression and improve clinical outcomes in HPV-associated cancers.

## Introduction

High-risk human papillomaviruses (HR HPVs) are etiologically linked to epithelial cancers, including ano-genital cancers and head and neck squamous cell carcinoma (HNSCC) (1). HPV is associated with around 80% of oropharyngeal cancers (HPV+OPC); the majority of patients with HPV+ OPC respond better to radiotherapy (2–5). Previous work has shown that, whereas the viral genome is found to be integrated into host DNA in the majority of cervical cancers, HPV DNA is episomal in around 70% of HPV+HNSCC (6, 7). The virus requires the differentiation program of epithelia in order to complete its life cycle, and utilizes host DDR proteins to replicate its genome (8, 9). In order to facilitate this, viral proteins E1 and E2 recruit host DDR factors to viral DNA; an active DDR is required for HPV life cycles (9–12). There are several components of the DDR pathway which facilitate the HPV life cycles; functional interactions occur between HPV16 and DDR factors including TOPBP1, SIRT1, WRN, BRD4 and SAMHD1 among others (13–19). Activation of the DDR by HPV16 promotes homologous recombination, which facilitates replication of the viral genome (20). During replication stress that occurs on the viral genome during the viral life cycle, fork reversal likely occurs and several host factors maintain these forks and resolve the reversal to continue ongoing replication (21, 22). One factor required for fork reversal is SMARCAL1 (23, 24).

SMARCAL1 is a member of the highly conserved SNF2 ATP-dependent chromatin-remodeling enzyme family (23, 25, 26). This family functions in gene transcription, DNA damage repair, DNA recombination, DNA methylation, and cell cycle regulation. Interaction of SMARCAL1 with RPA, via the RPA32 subunit, at stalled replication forks recruits SMARCAL1 to sites of replication stress where it catalyzes the rewinding of single-stranded DNA (27–29). SMARCAL1 is recruited to adenovirus viral replication centers (30). Given the role of SMARCAL1 in fork reversal, and previous involvement in DNA virus replication, we investigated the role of SMARCAL1 in the HPV16 life cycle.

Here we demonstrate that SMARCAL1 is recruited to HPV16 E1-E2 transiently replicating DNA in the non-HPV cervical cancer cell line C33a. Removal of SMARCAL1 in C33a cells using shRNA does not alter the levels of E1-E2 mediated DNA replication, but does reduce the fidelity of replication. In HPV16 immortalized human foreskin keratinocytes (HFK+HPV16) SMARCAL1 is more recruited to host replication forks when compared with control cells. We recently demonstrated that E2 can induce the DNA damage response (DDR) during mitosis and this could be contributory to DDR activation in the full HPV16 genome cells (31). SMARCAL1 is also recruited to the viral genome in HFK+HPV16 cells. Knockdown of SMARCAL1 in HFK+HPV16 attenuates cell growth; this does not occur in HFK expressing only the viral oncogenes E6 and E7. The HFK+HPV16 cells with SMARCAL1 knockdown have slower host replication fork progression, increased DNA damage, and integrated viral genomes. The results present a model in which the presence of HPV16 promotes recruitment of SMARCAL1 to viral and host DNA for correct viral and host replication fork progression. In the absence of SMARCAL1, the presence of HPV16 results in attenuation of host replication fork progression, increased double stranded DNA breaks (likely due to replication fork collapse) and the abrogation of cellular growth. Overall, the results present SMARCAL1 function as a therapeutic target for disrupting the growth of HPV16 positive cells, including cancers that retain episomal viral genomes.

## Results

### SMARCAL1 interacts with the viral replication complex and regulates viral replication fidelity

The recruitment of DDR factors to HPV replication centers demonstrates replication stress occurring on the viral genome. Our prior work demonstrated that WRN is critical for viral replication, suggesting that the process of replication fork reversal is critical for viral replication (17). To investigate whether SMARCAL1, a host protein also involved in replication fork reversal, is recruited to E1-E2 replicating DNA C33a cells were transiently transfected with plasmids containing the HPV16 origin of replication (pOri) and expressing the viral replication factors E1 and E2. Chromatin was isolated 48 hours following transfection and ChIP performed using antibodies to detect SMARCAL1, E1 (HA as the E1 expression vector is HA-tagged) and E2; all three factors were recruited to the E1-E2 replicating DNA (Figure 1A, lanes 10-12). Lanes 1-3 were transfected only with the plasmid containing the HPV16 origin (pOri), lanes 4-6 with pOri plus an E1 expression vector, and lanes 8-9 with pOri plus an E2 expression vector. Our previous work has demonstrated both E1 and E2 expression are required for viral factor recruitment which in turn recruits host factors (32).

**Figure 1.**
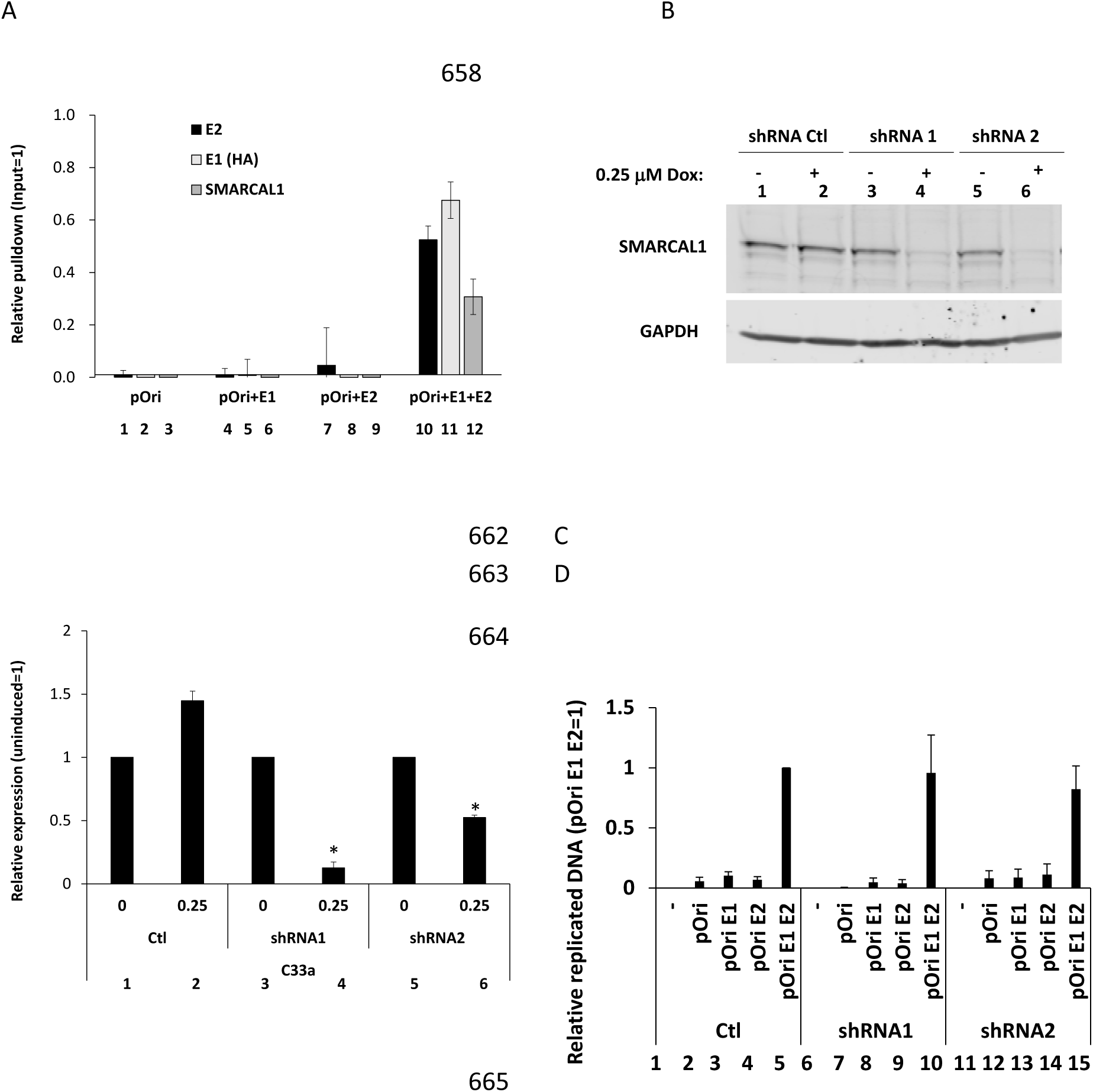
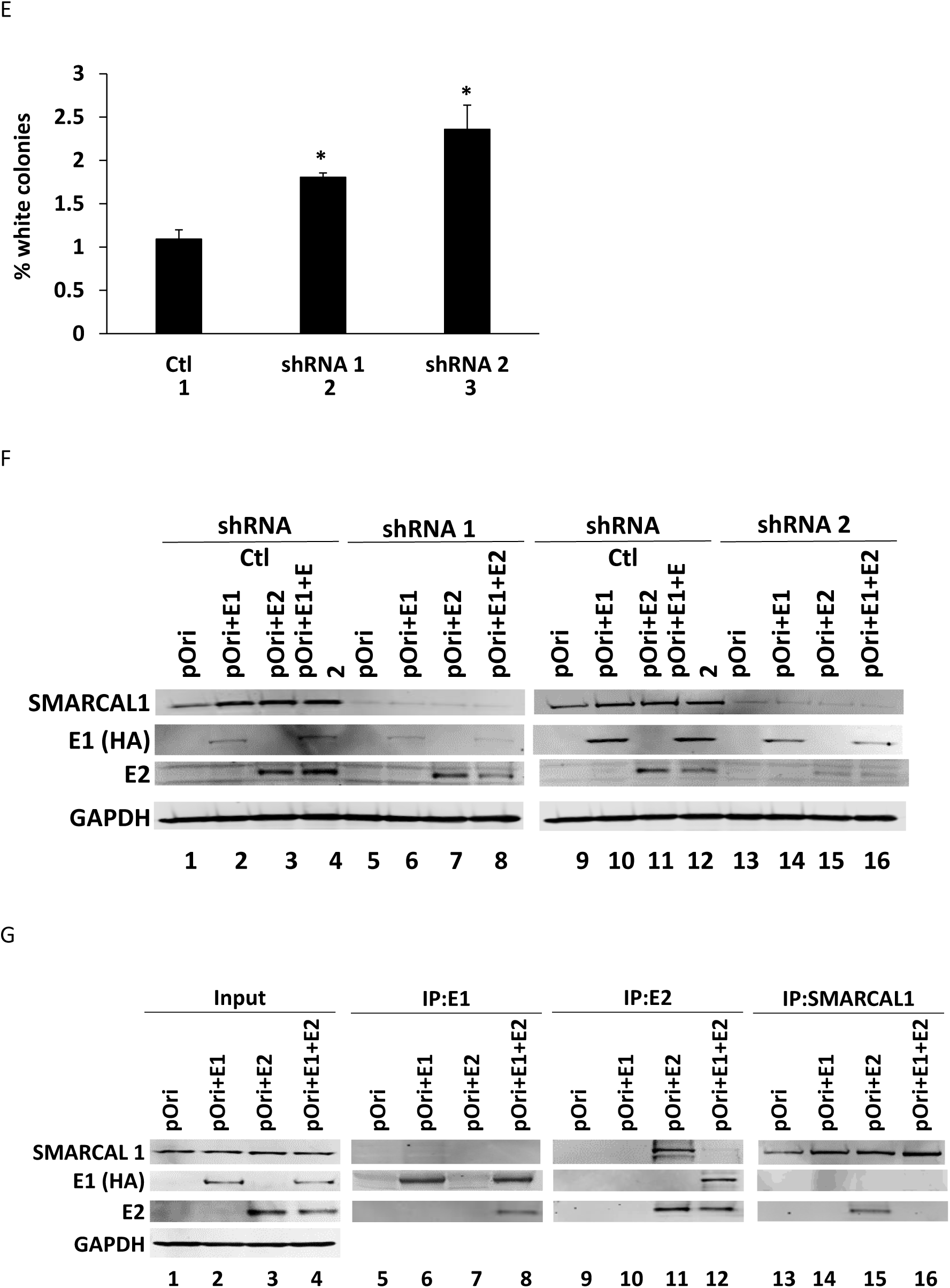
SMARCAL1 is recruited to E1-E2 replicating DNA. A. C33a cells were transfected with 1μg of pOri (a plasmid containing the HPV16 origin of replication), 1μg of an HPV16 E1 expression vector (HA tagged), and 1μg of an HPV16 E2 expression vector. 48-72 hours following transfection chromatin was prepared and ChIP assays carried out using E2, HA (for E1) and SMARCAL1 antibodies. The results are representative of a summary of three independent experiments. With all antibodies, there is a significant increase in pOri signal when E1 and E2 are expressed, versus no viral protein or either by itself. B. C33a cells expressing doxycycline (Dox) inducible control shRNA (shRNA Ctl) or two targeting SMARCAL1 (shRNA1 and shRNA2) were generated and 0.25mM of Dox added to confirm SMARCAL1 knockdown with the targeting shRNAs. C. This confirms SMARCAL1 RNA knockdown in the C33a cells represented in B. D. C33a cells were transfected with 1μg of pOri, 1μg of an HA-E1 expression vector, and 1μg of E2 expression vector. 48-72 hours following transfection cells were harvested and processed for DNA replication determination as described in materials and methods. The results presented represent the summary of at least three independent experiments. E. The DNA harvested in D was electroporated into bacteria and the number of white colonies determined on agar-x-gal plates. The number of white colonies identifies unfaithful replication. F. C33a cells in B/C were treated for 48 hours with 0.25 mM Dox and transfected with 1μg of pOri, 1μg of an HA-E1 expression vector, and 1μg of E2 expression vector. Protein extracts were prepared 48-72 hours later and western blotting carried out as indicated. G. C33a cells were transfected with 1μg of pOri, 1μg of an HA-E1 expression vector, and 1μg of E2 expression vector and proteins extracted 48-72 hours later. The indicated immunoprecipitations followed by western blotting are shown. * = p-value < 0.05.

To assess the functional impact of SMARCAL1 recruitment to E1-E2 replicating DNA, we generated C33a cell lines with inducible shRNA targeting SMARCAL1. Western blot analysis confirmed the knockdown of SMARCAL1 protein expression with two separate shRNAs following incubation of cells with 0.25 μM doxycycline for 48 hours (Figure 1B, lanes 4 and 6). qRTPCR also confirmed knockdown of SMARCAL1 RNA (Figure 1 C, lanes 4 and 6). Cell lines containing inducible control shRNA showed no decrease in SMARCAL1 protein or RNA following doxycycline treatment (Figure 1 B and C, compare lanes 1 and 2). Following induction of shRNA, cells were transfected with E1, E2 expression vectors and pOri and DNA harvested 48 hours later and processed for measurement of E1-E2 mediated DNA replication levels (32). Knockdown of SMARCAL1 did not significantly alter E1-E2 mediated DNA replication (Figure 1D, compare lanes 10 and 15 with lane 5). pOri contains the bacterial lacz gene and we have used this to determine replication fidelity; the percentage of white colonies signals mutations in the lacz gene introduced during E1-E2 replication (33). While the replication assays showed no significant change in levels following SMARCAL1 knockdown, mutation frequency increased significantly, indicating reduced viral replication fidelity in the absence of SMARCAL1 (Figure 1E, compare lanes 2 and 3 with lane 1). Knockdown of SMARCAL1 did not significantly alter expression of the E1 and E2 proteins in these C33a experiments (Figure 1F).

To investigate whether the viral replication factors could directly interact with SMARCAL1 immunoprecipitation (IP) experiments were performed (Figure 1G). Input levels of proteins for the IPs are shown lanes 1-4, E1 (HA) in lanes 5-8, E2 in lanes 9-12 and SMARCAL1 in lanes 13-16. E1 is unable to co-IP SMARCAL1 (lane 6) but as expected interacted with E2 (lane 8). E2 could co-IP SMARCAL1 when expressed by itself (lane 11), but was unable to do so when E1 is co-expressed (lane 12); E1 interacts with E2 as expected (lane 12). SMARCAL1 can interact with E2 (lane 15), but not when E1 is co-expressed (lane 16). This presents a model where E2 may be involved in recruiting SMARCAL1 to viral DNA but following E1 complexing with E2 SMARCAL1 is removed from E2. This is presumably due to a high affinity interaction between E1 and E2 displacing SMARCAL1 from E2. This mechanism would promote recruitment of SMARCAL1 to the viral genome.

### SMARCAL1 is essential for HPV-driven cell growth

Figure 1 demonstrates the involvement of SMARCAL1 in the regulation of E1-E2 mediated DNA replication in C33a cells, prompting investigation of the role of SMARCAL1 in the HPV16 life cycle. ChIP assays were carried out with chromatin isolated from keratinocytes immortalized with the full HPV16 genome (HFK+HPV16), utilizing SMARCAL1 antibodies, with HA and E2 antibodies as negative and positive controls (the E1 in the HPV16 genome is not tagged with HA, as it is in the C33a experiments carried out in Figure 1), respectively. In two donor keratinocyte backgrounds, SMARCAL1 was precipitated with viral DNA at a comparable level to viral protein E2 (Figure 2A, compare lane 1 with 2 and 3 with 4). Both SMARCAL1 and E2 interacted with the viral genome at significantly greater levels than the HA control antibody (lanes 5 and 6).

**Figure 2.**
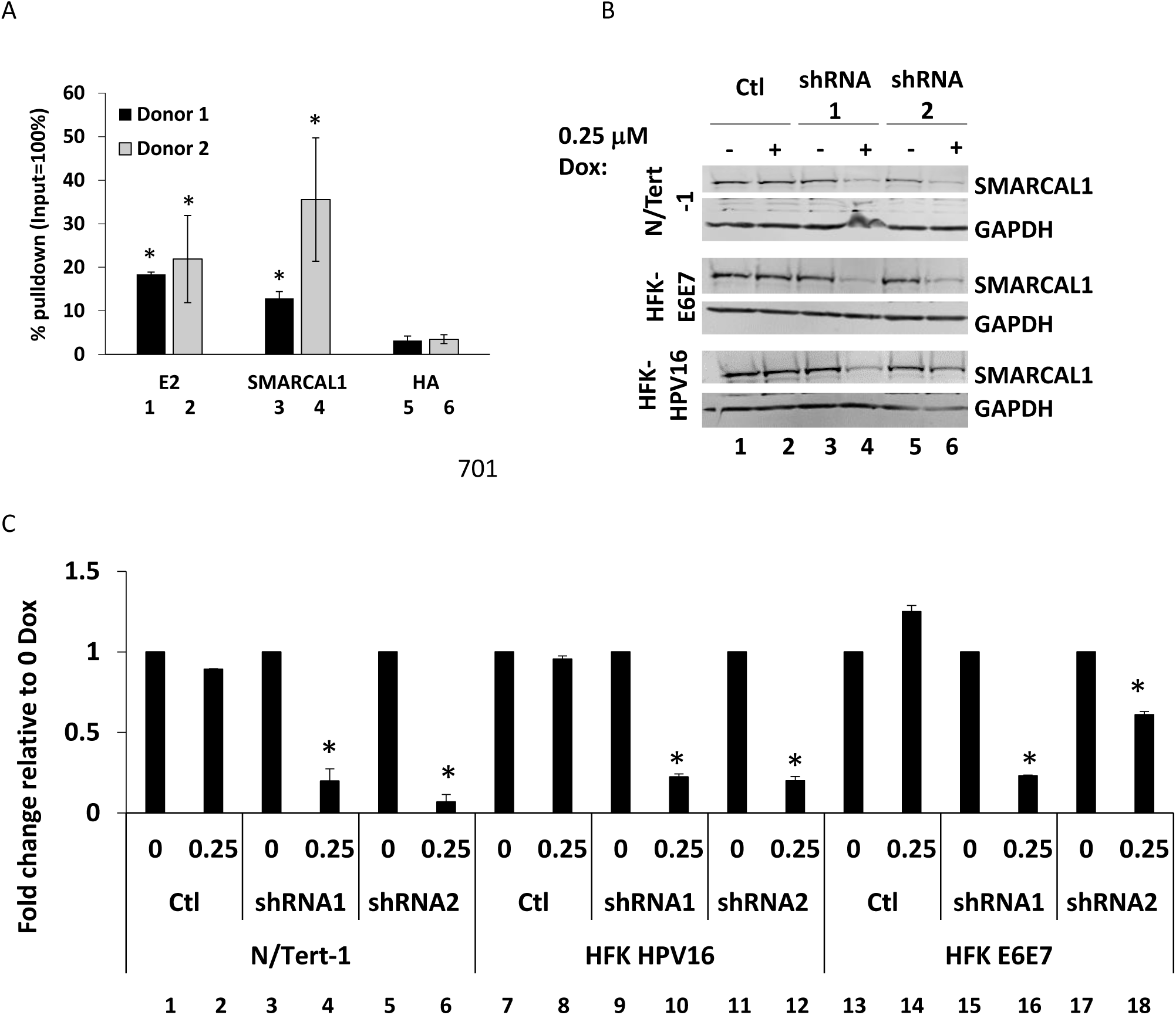
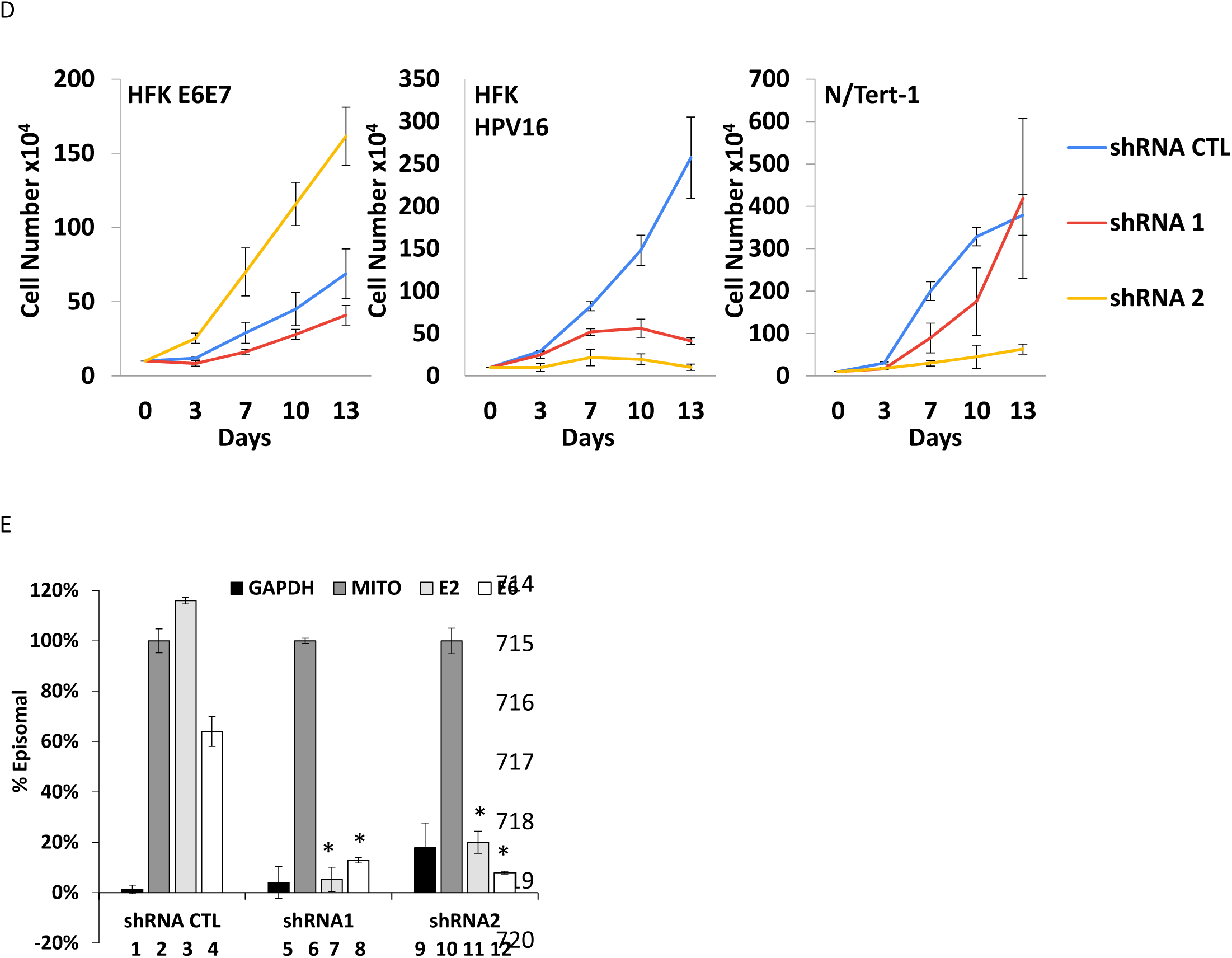
SMARCAL1 is critical for the growth of HFK+HPV16 cells. A. Chromatin was prepared from two HFK+HPV16 (human foreskin keratinocytes immortalized with HPV16) cell lines generated from two different donors and immunoprecipitated (ChIP) with E2, SMARCAL1 and HA (as a negative control) antibodies and the presence of the viral origin detected using qPCR. The results represent the summary of at least two independent experiments. B. N/Tert-1+Vec (containing pcDNA as a control), HFK+E6E7 (immortalized with the viral oncogenes E6 and E7 only) and HFK+HPV16 were generated expressing inducible shRNA control (Ctl) or inducible shRNAs targeting SMARCAL1 (shRNA1 and shRNA2). The cells were treated with 0.25μM Dox for 48 hours prior to protein harvesting, and western blotting confirmed knockdown of SMARCAL1 in the appropriate samples. C. Knockdown of SMARCAL1 RNA via the shRNA targeting sequences was confirmed. D. Growth curves of the indicated cell lines were carried out over the period indicated, cells were trypsinized and passaged at each of the days indicated. The results shown represent the summary of three independent experiments. E. Viral genome status was investigated using TV exonuclease assays following knockdown of SMARCAL1 and the episomal status of the E2 and E6 signal determined relative to that of mitochondrial DNA, which, like the viral genome is episomal. * = p-value < 0.05.

To investigate the role of SMARCAL1 in the HPV16 life cycle, HFK+HPV16 lines were established expressing either vector control (Ctl) or doxycycline-inducible shRNAs targeting SMARCAL1 (shRNA1-3). This was done in HFKs immortalized with either HPV16 full genome (HFK+HPV16) or the oncoproteins E6E7 (HFK+E6E7), or telomerase (N/Tert-1); the HFK+HPV16 and HFK+E6E7 were isogenic. shRNA expression was induced by culture in 0.25μM doxycycline and western blotting confirmed reduced SMARCAL1 protein expression following shRNA targeting of SMARCAL1, when compared with control shRNA (Figure 2B, compare lanes 4 and 6 with lane 2). qRT-PCR confirmed significant RNA knockdown of SMARCAL1 with the targeting shRNAs when compared with control shRNA (Figure 2C). To determine the effect of SMARCAL1 knockdown on cell growth, cells were counted over a 13-day period with passaging at days 3, 7 and 10 (Figure 2D). There was a variable effect on cell growth in N/Tert-1 and HFK+E6E7 cells following SMARCAL1 knockdown (Figure 2D, left and right panels). However, in HFK+HPV16 cells SMARCAL1 knockdown ultimately resulted in a growth stop (Figure 2D, middle panel). SMARCAL1 knockdown using both targeting shRNAs resulted in a significant reduction in growth of HFK+HPV16 cells.

Given the recruitment of SMARCAL1 to replicating HPV16 DNA in HFK+HPV16 cells (Figure 2A), and the role of SMARCAL1 in managing DNA replication stress, the genome status of HPV16 was determined following SMARCAL1 knockdown. The most quantitative assay for determining HPV16 genome status was recently demonstrated to be the TV exonuclease assay as it can measure the levels of both integrated and episomal genomes and was the most reflective of DNA-seq data (34). This approach was used to investigate the HPV16 genome status following SMARCAL1 knockdown. Mitochondrial DNA (which is circular, therefore episomal) was used as a control for episomal integrity and set as 100%. The signal generated with E2 demonstrates the E2 gene is on viral episomes (Figure 2E, lane 3); it is slightly over 100% as the mitochondrial DNA samples had, on average, a small amount of degradation in the DNA samples. E6 has less integrity but remains over 50% episomal (Figure 2E, lane 4), indicating that there is likely a mix of episomal and integrated genomes in the samples, with the integrated samples having lost the E2 gene. Following SMARCAL1 knockdown, the E2 and E6 signals become mostly integrated as the signals detected are similar to those of GAPDH (Figure 2E, lanes 7&8, and 11&12) and are significantly less than the levels detected with the control shRNA (lanes 3 and 4). It can be concluded from these studies that SMARCAL1 expression is essential for the maintenance of episomal viral genomes and is therefore essential for the HPV16 life cycle.

### SMARCAL1 is hyper recruited to host replication forks in HPV16+ keratinocytes

The essential nature of SMARCAL1 for the growth of HFK+HPV16 cells (Figure 2) prompted us to analyze the recruitment of SMARCAL1 to host replication forks. To investigate this, we performed SIRF assays (i*n <u>s</u>itu* analysis of protein *i*nteractions at DNA *<u>r</u>*eplication *f*orks). SIRF combines proximity ligation assays (PLA) with EdU click chemistry to detect protein co-localization with nascent DNA at a single cell level (35). To allow direct comparison within isogenic cell lines, we used our previously characterized N/Tert-1+Vec (vector control), N/Tert-1+HPV16 (containing the entire HPV16 genome), N/Tert-1+E6E7 (expressing only the E6 and E7 oncogenes) and N/Tert-1+HPV16 ΔE6/E7 in which stop codons were introduced in the E6 and E7 oncogenes were used; HPV16 ΔE6/E7 serves as control for the role of episomal replication without any effect of viral oncogene expression (36–38). The presence of HPV16 increased SMARCAL1 association with host replication forks. Figure 3A demonstrates that there is an increased presence of SMARCAL1 at host replication forks in N/Tert-1+HPV16 cells when compared with N/Tert-1+Vec as demonstrated by the increased number of red dots observed (representative images to the left, quantitation to the right; compare lanes 1 and 2). SMARCAL1 was recruited to host replication forks in N/Tert-1+E6E7 cells at significantly higher levels when compared with N/Tert-1+HPV16 (Figure 3A, compare lanes 2 and 3). SMARCAL1 was also more recruited to host replication forks in N/Tert-1+HPV16 ΔE6/E7 cells when compared with N/Tert-1+Vec control cells (Figure 3A, compare lanes 1 and 4). This suggests that both viral replication *per se* and the viral oncogenes can regulate SMARCAL1 recruitment to host replication forks. Interestingly, the SMARCAL1 recruitment in N/Tert-1+HPV16 and N/Tert-1+HPV16 ΔE6/E7 cells is not significantly different (compare lanes 2 and 4), suggesting that when the entire viral genome is present the regulation of SMARCAL1 recruitment is controlled by viral replication rather than the viral oncogenes. We will speculate further on the mechanisms of SMARCAL1 recruitment to host DNA replication forks by HPV16 in the discussion. The increased recruitment of SMARCAL1 by the viral oncogenes by themselves was reproducible in HFK+HPV16 versus HFK+E6E7 cells (Figure 3B, compare lanes 1 and 2 in the right-hand panel). Following replication stress induced by hydroxyurea, SMARCAL1 was recruited to the host forks in all N/Tert-1 cell lines at comparable levels demonstrating the capacity of these cells to recruit SMARCAL1 to host replication forks under replication stress conditions (see Supplementary Figure 1).

**Figure 3.**
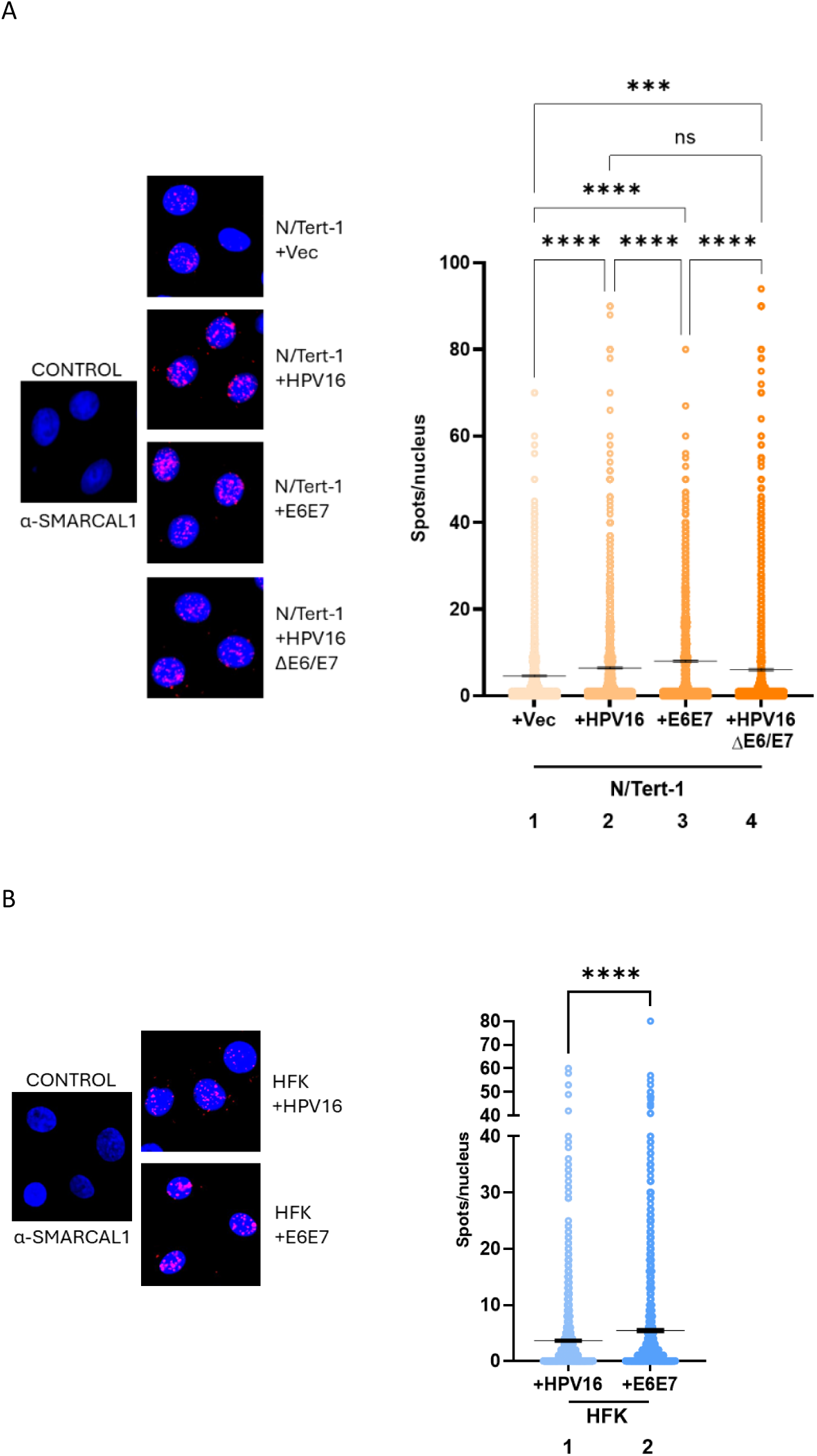
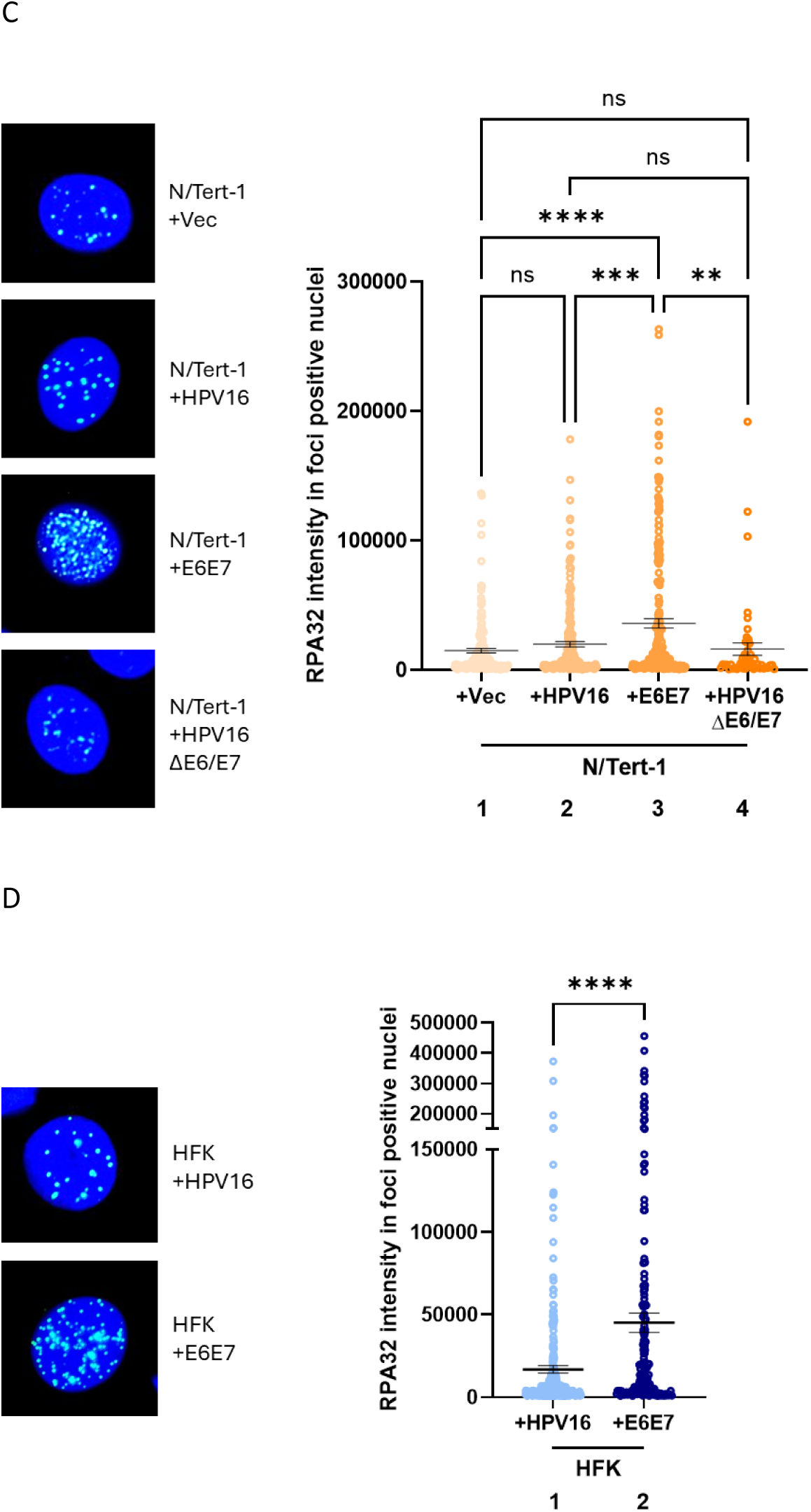
Elevated recruitment of SMARCAL1 in cells containing HPV16 or the viral oncogenes E6E7. A. SIRF assays were carried out with indicated cell lines and representative images are shown on the left. The controls represent SMARCAL1 only antibody on N/Tert-1+Vec lines. B. SIRF assays were repeated in HFK+HPV16 and HFK+E6E7 cell lines and representative images are presented on the left, and quantitation on the right. C. RPA32-foci staining was analysed by IF. The graphs show the intensity of RPA32-foci positive N/tert-1 (A) and HKF (B) cells measured from two independent experiments (at least 50 nuclei). On the left the representative images of RPA32-foci. The levels of statistical significance are indicated as P values of <0.01 (**), <0.001 (***) and <0.0001 (****); ns, not significant (One-way Anova test for panel B, Mann-Whitney test for panel D).

Exogenous expression of the viral oncogenes by themselves can induce replication stress (39), to investigate the levels of replication stress in the cell lines under study, immunofluorescence assays were carried out with RPA32, a known readout of replication stress (40). Figure 3C demonstrates that there is not an increase in replication stress in N/Tert-1+HPV16 when compared with N/Tert-1+Vec control cells (compare lanes 1 and 2). However, cell lines overexpressing the viral oncogenes E6E7 by themselves demonstrate a significantly increased level of DNA replication stress when compared with all other N/Tert-1 cell lines (compare lane 3 with lanes 1, 2 and 4). Similarly, in HFK+E6E7 cells there is a significant increase in replication stress when compared with HFK+HPV16 (Figure 3D, compare lanes 1 and 2). These results suggest that the enhanced recruitment of SMARCAL1 to host replication forks in cells containing the entire HPV16 genome may not be due to simple replication stress, but is due to an alternative mechanism.

Overall, these results demonstrate that both replication of the viral genome, and the viral oncogenes by themselves, recruit SMARCAL1 to host replication forks for fork reversal resolution and ongoing replication. Our prior work demonstrated that N/Tert-1+HPV16 ΔE6/E7 cells retained an activate DDR providing a potential mechanism for the recruitment of SMARCAL1 to host replication forks in these cells (38).

### Knockdown of SMARCAL1 increases DNA damage in HFK+HPV16 cells

The results so far have determined that SMARCAL1 is recruited to HPV16 E1-E2 replicating DNA (Figures 1 and 2), that SMARCAL1 is essential for the growth of HFK+HPV16 cells (Figure 2), and that SMARCAL1 is hyper-recruited to host replication forks in the presence of the HPV16 genome and the viral oncogenes (Figure 3). The reason for the essential nature of SMARCAL1 in the growth of HFK+HPV16 was investigated. Figure 4A quantitates a COMET assay in N/Tert-1+Vec, HFK+HPV16 and HFK+E6E7 cells with and without SMARCAL1 expression. In shRNA control (shRNA CTL) cells HFK+HPV16 have significantly longer olive tail moments (OTMs) when compared with N/Tert-1+Vec and HFK+E6E7 (compare lane 7 with lanes 1 and 4, respectively) indicative of increased DNA damage. When SMARCAL1 is knocked down by shRNA1&2 in HFK+HPV16 cells there is a significant increase in DNA damage when compared with HFK+HPV16 shRNA CTL (compare lanes 8 and 9 with lane 7). This increased DNA damage correlates with the critical nature of SMARCAL1 for HFK+HPV16 cell growth (Figure 2). There is not a consistently significant increase in OTM length in either N/Tert-1+Vec or HFK+E6E7 following SMARCAL1 knockdown (compare lanes 2 and 3 with lane 1, and lanes 5 and 6 with lane 4, respectively). As SMARCAL1 knockdown cannot consistently abrogate the growth of either of these cell lines, this further supports the idea that in HFK+HPV16 cells SMARCAL1 knockdown results in growth attenuation due to increased DNA damage.

**Figure 4.**
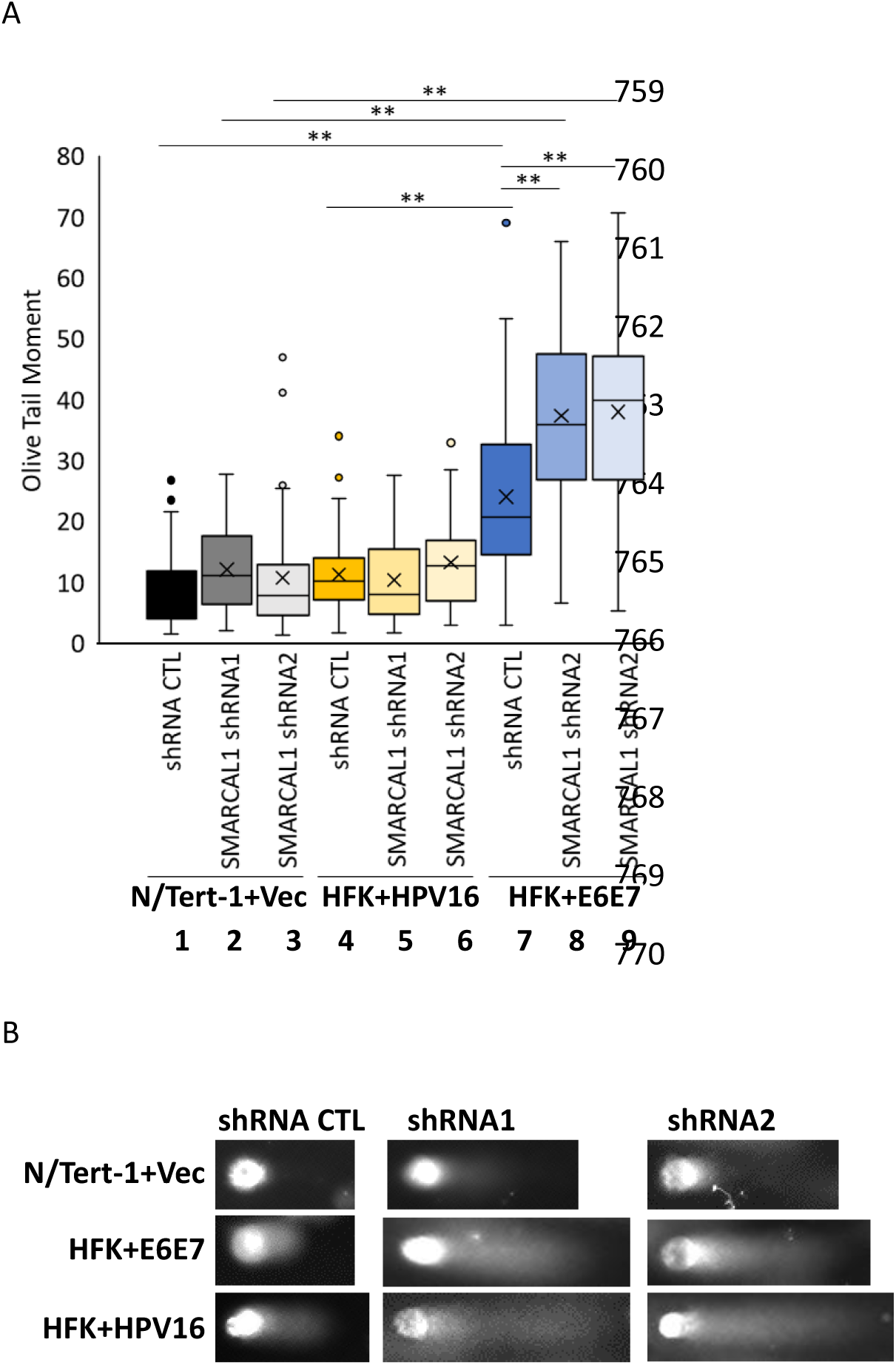
SMARCAL1 reduction induces DNA damage in HFK+HPV16 cells. A. COMET assays were carried out with the indicated cell lines 48 hours following treatment with 0.25μM Dox. A summary of the olive tail moments is shown. B. Representative DNA “tails” in the indicated cell types. ** = p-value < 0.05.

To determine whether HFK+HPV16 increased DNA damage in the absence of SMARCAL1 was the result of host replication forks slowing down (ultimately resulting in double strand breaks), fiber assays were utilized (Figure 5). The quantitated results are shown in Figure 5A. In HFK+HPV16 cells SMARCAL1 knockdown significantly slowed replication fork speed (compare lane 7 with lanes 8&9), and the slowdown in fork speed was significantly more than that observed in HFK+E6E7 cells (lanes 5 and 6), or in N/Tert-1+Vec cells (lanes 2 and 3). SMARCAL1 knockdown did decrease fork speed in HFK+E6E7 cells (compare lanes 5 and 6 with lane 4), and was variable in N/Tert-1 cells (compare lanes 2 and 3 with lane 1). Overall, the results demonstrate that SMARCAL1 knockdown had the greatest effect on fork speed in HFK+HPV16 cells and this correlates with the increased double strand DNA breaks detected in these cells (Figure 4).

**Figure 5.**
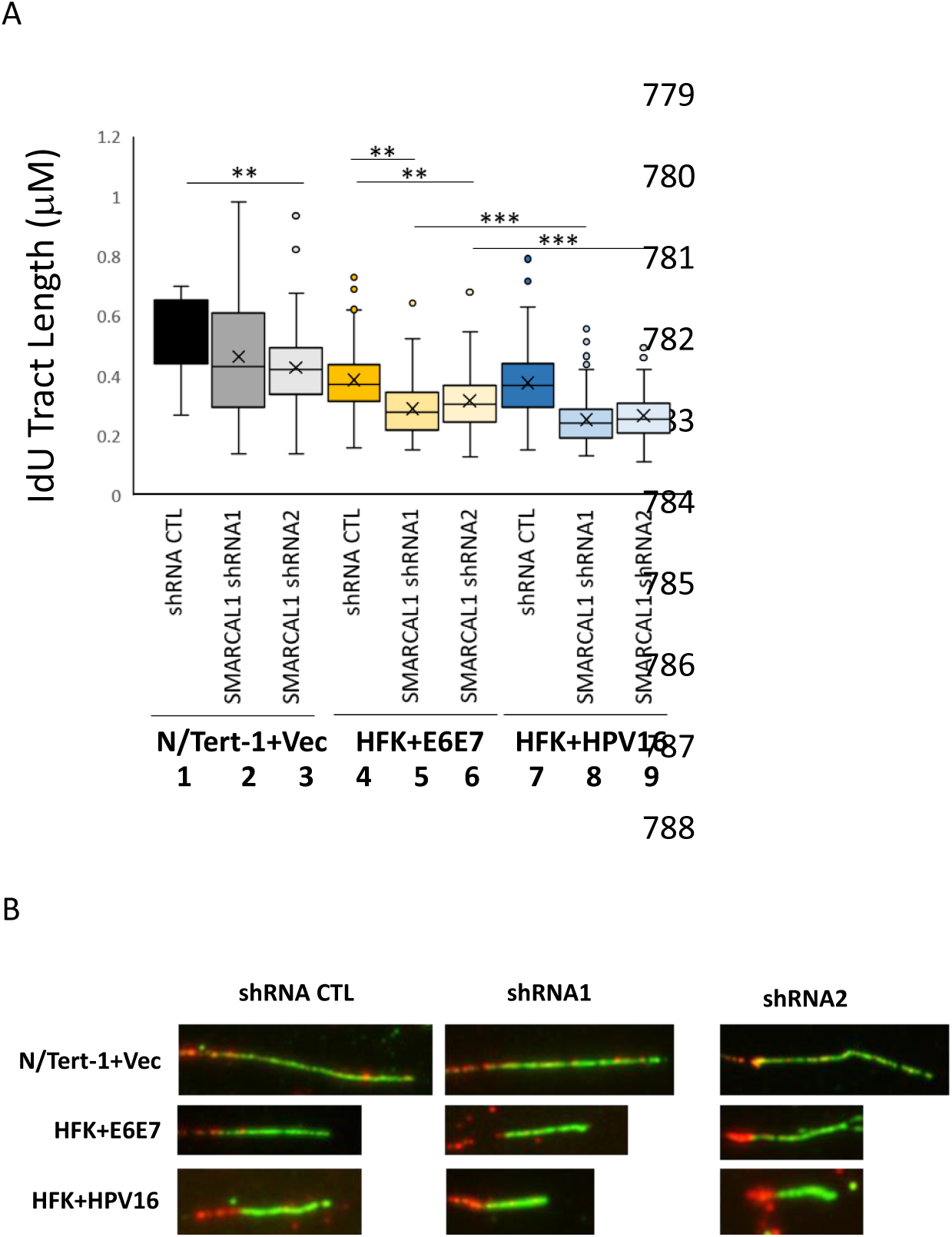
SMARCAL1 reduction attenuates host fork replication speeds in HFK+HPV16 cells. A. Fiber assays were carried out with the indicated cell lines 48 hours following treatment with 0.25μM Dox. A summary of the “green” fiber length is shown. B. Representative fibers from the indicated cell types. The * highlight differences between samples, and the larger the number of * the higher the significance. * = p-value < 0.05 in all cases.

## Discussion

It is well established that HPV life cycles require activation of the DNA damage response (DDR) during epithelial differentiation (9). The reason for this activation is presumably to facilitate viral replication which occurs more than once per cell cycle at early and late stages of the viral life cycle, with the potential to generate replication stress on the viral genome. Activation of the DDR promotes activation of several homologous recombination (HR) factors that would promote viral genome replication. Our previous work demonstrated that the host protein WRN is critical for E1-E2 DNA replication and for the viral life cycle (17). WRN is actively involved in several DNA repair pathways, including the resolution of replication fork reversals that can occur under replication stress conditions (41, 42). Given the critical role of WRN in the viral life cycle and its known involvement in stalled replication fork reversal resolution, we investigated the role of another protein involved in replication fork reversal resolution, SMARCAL1, in the viral life cycle (24, 25, 27, 28). The results presented here demonstrate that SMARCAL1 is recruited to E1-E2 replicating DNA and is required for maintaining high fidelity E1-E2 replication. Importantly, depletion of SMARCAL1 results in viral genome integration, therefore SMARCAL1 is essential for the HPV16 life cycle as it is required for viral genome maintenance. Our understanding of mechanisms and factors involved in maintaining an episomal viral genome is incomplete, and the results presented here demonstrate SMARCAL1 is critical for viral genome maintenance. Therefore, SMARCAL1 can be added to the growing list of DNA damage/repair factors that are important during the viral life cycle. An interesting aspect of our results is that elimination of SMARCAL1 from cells preferentially attenuates the growth of keratinocytes containing the entire HPV16 genome. Many studies have demonstrated host factors are important for viral genome maintenance as their knockdown promotes viral genome integration, including SIRT1 and members of the MRN complex, but that the presence of these factors is not essential for the growth of cells with entire HPV genomes (19, 43–45). The essential nature of SMARCAL1 for HPV16 positive cell growth (but not those immortalized by the viral oncogenes E6 and E7) prompted us to investigate the mechanism of why SMARCAL1 depletion preferentially abrogates growth in cells containing the full HPV16 genomes.

Using SIRF assays we demonstrate that SMARCAL1 is recruited to host DNA replication forks in cells containing the entire HPV16 genome, or the viral oncogenes E6 and E7 only. Also, a viral genome with stop codons in E6 and E7 recruits SMARCAL1 to host replication forks to the same extent as cells containing the full genome. The viral oncogenes by themselves are “better” promoters of SMARCAL1 localization to host replication forks. What is the explanation for these differences? We propose that in cells containing the viral oncogenes only, the replication stress induced (as demonstrated by the elevated presence of RPA at host replication forks) promotes recruitment of SMARCAL1 to the host forks. It is noticeable that replication stress, as measured by RPA recruitment, is not increased in cells containing the entire HPV16 genome, but SMARCAL1 is still increased in recruitment to host DNA replication forks. Recently, we demonstrated that the presence of E2 (present in the HPV16 cells, not present in the oncogene only cells) induces an active DDR during mitosis (we have extended these studies to demonstrate E2 activates the DDR during differentiation, not shown) (31). We propose that this mitotic DDR promotes damage on the host DNA and this is supported by the increased COMET tails detected in cells containing HPV16 versus the E6 and E7 oncogenes only. This increased damage may alter the requirements at host replication forks during the subsequent S phase promoting recruitment of SMARCAL1, which is subsequently required to limit further accumulation of DNA damage. If this is the driver for SMARCAL1 recruitment in cells containing the full HPV16 genome, it would also explain why SMARCAL1 has increased recruitment to host replication forks in HPV16 ΔE6/E7 cells. Therefore, there are two different mechanisms the virus uses to induce SMARCAL1 to host replication forks, one related to viral replication (perhaps E2 expression) and one related to the presence of the viral oncogenes. The reason that the two different mechanisms are not additive in cells with the entire HPV16 genome could be that the viral oncogenes are not as “active” in cells containing the entire genome, as traditional E6 and E7 targets, p53 and pRB respectively, are expressed in cells containing the entire HPV16 genome, but not in cells immortalized only with the viral oncogenes (46). We are currently investigating the reasons for the differences in oncogene function under different circumstances, and further elucidating the mechanism of SMARCAL1 recruitment to host DNA replication forks by HPV16.

SMARCAL1 is also critical for viral replication as it is recruited to E1-E2 replicating DNA, and depletion of SMARCAL1 results in a reduction in E1-E2 replication fidelity. How SMARCAL1 is recruited to the viral replication forks is unclear, it could be related directly to the DNA structures generated on the viral genome. But, it is also of interest that E2 can interact with SMARCAL1 and it is possible that E2 recruits SMARCAL1 to the viral replication origin. The E2-SMARCAL1 interaction is disrupted in the presence of E1 suggesting that E1 and SMARCAL1 interact with E2 in similar areas. Interestingly, WRN interacts with E1 in the domain of E1 that interacts with E2 (17). This suggests an intricate interaction of the viral replication factors with host DDR proteins that is required for viral replication and execution of the viral life cycle.

Overall, the results demonstrate a critical role for SMARCAL1 in the viral life cycle. The results also present a novel approach to the targeting of HPV16 infected cells; inhibition of SMARCAL1 enzyme function. Unlike other DNA damage factors involved in the viral life cycle, whose knockout induces viral genome integration but do not affect cell growth, SMARCAL1 depletion attenuates the growth of HPV16 positive keratinocytes. Future studies will focus on gaining an in depth understanding of why SMARCAL1 is essential for the growth of HPV16 positive cells, and developing SMARCAL1 as anti-viral therapeutic target.

## Materials and Methods

### Cell culture

C33a were obtained from ATCC (Manassas, VA, USA) and cultured in DMEM (Invitrogen) supplemented with 10% fetal bovine serum (FBS). Human foreskin keratinocytes (HFKs) were immortalized with HPV16 as described previously (47) or with 16 E6E7 by retroviral delivery using pLXSN16E6E7, a gift from Denise Galloway (Addgene plasmid #52394). HFKs were cultured in DermaLife-K Complete media (LifeLine Cell Technologies). Mitomycin C-treated 3T3-J2 fibroblasts feeders were plated 24 hours prior to seeding N/Tert-1 or HFK cells on top of the feeders, in their respective cell culture media. Media was refreshed and 3T3-J2s supplemented as required. N/Tert-1 cells were cultured in keratinocyte serum-free medium (K-SFM) (Invitrogen) supplemented with bovine pituitary extract, EGF (Invitrogen), 0.3 mM calcium chloride (MilliporeSigma; 21115) and 7.5 uM hygromycin. In all cases, cells were incubated at 37 °C in a 5% CO_2_/95% air atmosphere, routinely passaged before reaching confluency and screened for mycoplasma.

### Generation of inducible SMARCAL1 knockdown cell lines

A set of three SMARTvector^TM^ human Inducible Lentiviral shRNA plasmids targeting SMARCAL1 were purchased from Horizon, along with one empty vector control. C33a and keratinocytes immortalized with hTERT (N-Tert/1), E6E7 (HFK-E6E7) or HPV16 (HFK-HPV16) were infected with the resulting lentiviruses and selected with puromycin to generate lines expressing inducible control (CTL) or SMARCAL1 shRNA (shRNA1-3). Optimal doxycycline treatment for shRNA induction was determined by treatment of cells followed by PCR and western blot analysis; 0.25 uM doxycycline for 48 hours was found to induce shRNA without toxicity.

### Chromatin Immunoprecipitation (ChIP)

Cross-linking and chromatin extraction of C33a cells was carried out as previously described (ref). For chromatin extraction from keratinocytes, the ChIP-It Enzymatic kit (Active Motif) was utilized, as the protocol dictated, including dounce homogenisation to disrupt cells and incubation with shearing enzymes for 7.5 mins with frequent agitation. In both cases, sheared chromatin was incubated with 1 ug primary antibody (HA, Abcam ab9110; SMARCAL1 (Abcam ab154226); E2 Sheep) and protein-A conjugated magnetic beads, rotating overnight at 4°C. Beads were then washed and DNA eluted via Proteinase K digestion. Precipitated viral DNA was measured via qPCR using primers to the HPV16 LCR; Fwd 5’-GAAAACGAAAAGCTACACCCA-3’, Rev 5’-CAATGAATAACCACAACACAATTA-3’. Percentage pulldown was calculated by calculation to input: % Pulldown =100x 2^Input^ ^Ct-ChIP^ ^Ct^

### Transient Replication Assay

C33a cells were plated out at 5×10^5^ in 10 cm dishes. The following day, plasmid DNA was transfected using the calcium phosphate method (Kingston *et al.* 2003). Three days post-transfection, low molecular weight DNA was extracted using the Hirt method as previously described by Boner *at al*. 2002. The digested sample was extracted twice with phenol:chloroform:isoamyl alcohol (25:24:1) and precipitated with ethanol. Following centrifugation, the DNA pellet was washed with 70% ethanol, dried and resuspended in a total of 150 µL water. Forty-two µL of sample were digested with DpnI (New England Biolabs, Ipswich, MA, USA) overnight to remove unreplicated pOri16LacZ; the sample was then digested with ExoIII (New England Biolabs) for 1 hr. Replication was determined by real-time PCR, as described previously (Taylor *et al.* 2003).

### DNA Mutagenesis Assay

Replication fidelity was determined by blue white assay as described previously (Taylor *et al.* 2003). Essentially, DNA from the transient replication assay was re-suspended in 150µL of 10% glycerol; 75µL was electroporated into DH10B bacteria and plated onto kanamycin Lysogeny Broth (LB) agar containing 100 ug/mL X-gal. Following incubation overnight at 37°C, bacterial colonies were observed and counted; blue colonies correspond to those with no mutations in the LacZ gene, downstream of pOri, and the white colonies as those containing mutations. Thus, the ratio of blue colonies to white provides a readout of replication mutation frequency.

### In situ analysis of protein interactions at DNA replication forks (SIRF)

Exponentially growing cells were seeded onto microscope chamber slides. The day of the experiment, cells were incubated with 125 µM 5-ethynyl-2’-deoxyuridine (EdU) for 8 min. Cells were pre-extracted in 0.5% TritonX-100 for 10 min on ice and fixed with 2% PFA in PBS 1X for 15 min at room temperature. Cells were then permeabilized in 0.25% TritonX-100 for 15 min at RT. To detect EdU the Click-iT™ EdU Alexa Fluor™ Imaging Kit (Invitrogen) using 5 µM Biotin-Azide was used for 30 minutes at room temperature. Cells were washed with PBS 1X for 5 min and blocked in 3% BSA/PBS for 20 min. The primary antibodies used were as follows: SMARCAL1 (Abcam, 1:400), mouse anti-biotin (Jackson, 1:1200). The negative controls were obtained by using only one primary antibody. Samples were incubated with secondary antibodies conjugated with PLA (proximity ligation assay) probes MINUS and PLUS: the PLA probe anti-mouse PLUS and anti-rabbit MINUS (or NaveniFlex equivalent). The incubation with all antibodies was carried out in a humidified chamber for 1 hour at 37 °C. Next, the PLA probes MINUS and PLUS were ligated using two connecting oligonucleotides to produce a template for rolling-cycle amplification. After amplification, the products were hybridized with red fluorescence-labelled oligonucleotide. Samples were mounted in Prolong Gold anti-fade reagent with 4,6-diamidino-2-phenylindole (DAPI) (blue). Images were acquired randomly using an Eclipse 80i Nikon Fluorescence Microscope, equipped with a Virtual Confocal system. The analysis was carried out by counting the PLA spots for each nucleus, and plotted and analyzed using GraphPad Prism.

### Immunofluorescence (IF)

Cells were grown on coverslips in 35-mm dishes until 70-80% confluence. Subsequently, cells were washed with ice-cold PBS for 5 min, pre-extracted on ice with 0.5% Triton X-100 for 10 min and fixed with 3% para-formaldehyde (PFA)/2% sucrose at room temperature (RT) for 10 min. After two washes in PBS for 5 min, 3% bovine serum albumin (BSA) was added for 15 min at RT. Then, cells were incubated with anti-RPA34-20 (Merck, 1:400) diluted in a 1% BSA/0.1% saponin in PBS solution, for 1 h at 37 °C in a humidifier chamber. After three washes in PBS for 5 min, secondary antibody (Alexa Fluor 488-conjugated Goat Anti-Mouse IgG (H + L), highly cross-adsorbed; Life Technologies) was added at 1:200 dilution in a 1% BSA/0.1% saponin in PBS solution, for 1 h at RT. Counterstaining was performed with 0.5μg/ml DAPI. Slides were analysed with an Eclipse 80i Nikon Fluorescence Microscope, equipped with a Video Confocal (ViCo) system at 40× magnification. Fluorescence intensity for each sample was then analyzed using ImageJ software.

### Single-cell gel electrophoresis/Comet Assay

DNA breakage induction was evaluated by comet assay (single-cell gel electrophoresis) in alkaline conditions. Ten thousand cells were harvested by trypsinization and resuspended in 100µL of 1% (w/v) Seaplaque agarose (Lonza)/PBS. The cell-agarose mix was then transferred onto slides pre-treated with 1% agarose/PBS and left to set at 4°C in the dark. Slides were immersed in freshly prepared cold lysing solution (2.5M NaCl, 83 mM sodium-EDTA, 10mM Tris, 1% Triton X-100, 10% DMSO, pH10) and incubated for 1.5 hours at 4°C in the dark. Following this incubation, the slides were removed from the lysing solution and placed in a horizontal gel electrophoresis tank, side-by-side, with the agarose gels in the same plane of orientation in line with the anode. The electrophoresis tank was then filled with fresh electrophoresis buffer (0.3 M NaOH, 1 mM EDTA), and the slides were incubated in this buffer for 20 minutes. Cells were electrophoresed for 30 minutes at 20V. Following electrophoresis, slides were washed with neutralisation buffer (100 mM Tris pH 7.4) three times at 4°C, and stained for 3 minutes with 50 ul of 10 ug/ml ethidium bromide in distilled water. Excess stain was removed by flooding the slides with PBS and coverslips were applied prior to viewing using a fluorescence microscope (Keyence). DNA damage was quantified using CaspLab (kkoncr), 100 cells were analysed per sample, 50 cells per duplicate slide. The parameter used to reflect the amount of DNA damage was the Olive Relative Tail Moment (OTM), which is proportional to the % DNA in the tail multiplied by the distance between the two centers of mass in the head and tail (Olive et al.1995). This parameter was decided upon as the most appropriate descriptor of the fraction of DNA released from the cell nucleus and the distance migrated by this DNA in an electric field.

### DNA Fiber Analysis

Replication fork speed was measured using the DNA fiber assay. DNA fibers were prepared and analysed as described by Parra & Windle, 1993 and by Halliwell *et al.* 2020. First, cells were treated with 25 μM IdU for 20 minutes, and then cells were washed twice with PBS and incubated for 20 min in fresh medium containing 100 μM CldU. Cells were harvested by trypsinization and pellets resuspended in ice cold PBS. 2-4 μl of ice-cold cell suspension transferred to the top of each microscope slide, allowed to dry for 30 sec before adding 12 μL fresh lysis buffer (200mM Tris-HCl pH 7.5, 50 mM EDTA, 0.5% SDS) and stirring with the pipette tip. Slides were tilted to an angle of 25° to 40° for 2 minutes, and laid flat to air dry completely in the dark (approximately 5-15 min). Slides were fixed in fresh ethanol/acetic acid 3:1 for 15 minutes. Once dried, these were subject to immunodetection of labelled tracks, using the following primary antibodies were used: rat anti-CldU/BrdU (Abcam) and mouse anti-IdU/BrdU (Becton Dickinson). Images were acquired randomly from fields with untangled fibres using a Keyence BZ-X800 microscope. The length of labelled tracks was measured using the ImageJ software. A minimum of 100 individual fibres were analysed for each experiment and each experiment was repeated two times.

### Immunoblotting

Specified cells were trypsinized, washed with PBS and resuspended in 2x pellet volume NP40 protein lysis buffer (0.5% Nonidet P-40, 50 mM <u>Tris</u> [pH 7.8], 150 mM NaCl) supplemented with protease inhibitor (Roche Molecular Biochemicals) and phosphatase inhibitor cocktail (MilliporeSigma). Cell suspension was incubated on ice for 20 min and then centrifuged for 20 min at 184,000 rcf at 4 °C. Protein concentration was determined using the Bio-Rad protein estimation assay according to manufacturer’s instructions. 50 μg protein was mixed with 2x Laemmli sample buffer (Bio-Rad) and heated at 95 °C for 5 min. Protein samples were separated on <u>Novex</u> 4–12% Tris-glycine gel (Invitrogen) and transferred onto a nitrocellulose membrane (Bio-Rad) at 35V overnight using the wet-blot transfer method. Membranes were then blocked with Odyssey (PBS) blocking buffer diluted 1:1 with PBS and probed with indicated primary antibody diluted in Odyssey blocking buffer. Membranes were washed with PBS supplemented with 0.1% Tween (PBS-Tween) and probed with the Odyssey secondary antibody (goat anti-mouse IRdye 800CW or goat anti-rabbit IRdye 680CW) (Licor) diluted in Odyssey blocking buffer at 1:10,000. Membranes were washed twice with PBS-Tween and an additional wash with 1X PBS. After the washes, the membrane was imaged using the Odyssey^®^ CLx Imaging System and ImageJ was used for quantification, utilizing GAPDH as internal loading control. Primary antibodies used for western blotting studies are as follows: 16E2 monoclonal B9 1/500 (31), GAPDH 1/10,000 (Santa Cruz, sc-47724), HA-Tag (Abcam, ab9110), SMARCAL1 1/1000 (Abcam, ab154226).

### Immunoprecipitation (IP)

Cell lysate was prepared as described above. 500 µg of the lysate was incubated with lysis buffer (0.5% Nonidet P-40, 50mM Tris [pH 7.8], and 150mM NaCl), supplemented with protease inhibitor (Roche Molecular Biochemicals) and phosphatase inhibitor cocktail (MilliporeSigma) to a total volume of 500 µl. Primary antibody of interest or a FLAG-tag antibody (used as a negative control) was added to this prepared lysate and rotated at 4°C overnight. The following day, 40 µl of protein A beads per sample (MilliporeSigma; equilibrated to lysis buffer as mentioned in the manufacturer’s protocol) was added to the above mixture and rotated for another 4 hours at 4°C. The samples were gently washed with 500 µl lysis buffer by centrifugation at 1000 x g for 2-3 min. This wash was repeated 3 times. The bead pellet was resuspended in 4X Laemmli sample buffer (Bio-Rad), heat denatured and centrifuged at 1000 x g for 2-3 min. The supernatant applied to an SDS-PAGE system to separate and resolve proteins, and was then transferred onto a nitrocellulose membrane using wet-blot transfer method. The membrane was probed for the presence of as mentioned in the description of western blotting above.

### Exonuclease 5 (T5) Assay

PCR based analysis of viral genome status was performed using methods described by Myers *et al.* (2019). Briefly, 20 ng genomic DNA was either treated with exonuclease V (RecBCD, NEB), in a total volume of 30 ul, or left untreated for 1 hour at 37°C followed by heat inactivation at 95°C for 10 minutes. 2 ng of digested/undigested DNA was then quantified by real time PCR. Separate PCR reactions were performed to amplify HPV16 E6 F: 5’-TTGCTTTTCGGGATTTATGC-3’ R: 5’-CAGGACACAGTGGCTTTTGA-3’, HPV16 E2 F: 5’-TGGAAGTGCAGTTTGATGGA-3’ R: 5’-CCGCATGAACTTCCCATACT-3’, human mitochondrial DNA F: 5’-CAGGAGTAGGAGAGAGGGAGGTAAG-3’ R: 5’-TACCCATCATAATCGGAGGCTTTGG −3’, and human GAPDH DNA F: 5’-GGAGCGAGATCCCTCCAAAAT-3’ R: 5’-GGCTGTTGTCATACTTCTCATGG-3’. Episomal status was calculated by dCt(Exo-no Enzyme) and then comparison of dCt(HPV gene) to dCt(mitochondrial DNA) and dCt(GAPDH).

### Statistics

Standard error was calculated from three independent experiments and significance determined using a student’s t-test or Mann-Whitney for non parametric distribution. One-way Anova with Tukey’s test was used for multiple comparisons.

## Acknowledgements

This work was supported by US NIH grant R01DE029471 (IMM) and R21AI178143 (IMM). The funders had no role in study design, data collection and analysis, decision to publish, or preparation of the manuscript.

